# Bibliometric and Temporal Trend Analysis of Nipah Virus- An Emerging Zoonotic Disease: What Do We Know So Far

**DOI:** 10.1101/2023.10.17.562837

**Authors:** Rajeev K. Singla, Yingbo Zhang, Shailja Singla, Bairong Shen

**Affiliations:** Joint Laboratory of Artificial Intelligence for Critical Care Medicine, Department of Critical Care Medicine and Institutes for Systems Genetics, Frontiers Science Center for Disease-related Molecular Network, West China Hospital, Sichuan University, Chengdu, China; School of Pharmaceutical Sciences, Lovely Professional University, Phagwara, Punjab-144411, India; iGlobal Research and Publishing Foundation, New Delhi, India

**Keywords:** Nipah Virus, Henipavirus, Paramyxoviridae, Zoonotic Disease, Disease Outbreak

## Abstract

**Background:** Nipah virus (Genus Henipavirus) was first detected in 1999, during the Malaysia-Singapore outbreak, and is an emerging bat-borne pathogen. It causes rare but fatal disease, with a 40-75% fatality rate, and clinically ranging from asymptomatic infection to fatal encephalitis.

**Methods:** Publicly available literature, including preclinical and clinical studies, have been retrieved from PubMed, Scopus, and Web of Science. Bibliometric analysis was done using embedded tools of these search engines along with VOSviewer.

**Results:** 601 documents from PubMed, 955 from Web of Science, and 784 from Scopus were analyzed, and we found that maximum contributions are from the USA, India, Malaysia, Australia, and Bangladesh, and in the past five years, there has been an exponential surge in the publication rate. More research and high-throughput screening are needed to explore drugs against critical targets and receptors like ephrin-B2, non-structural protein C, F protein, L protein, G glycoprotein, nucleocapsid protein, V protein, P protein, and W protein. Research for possible vaccination is underway, but the rate is not significant. Clinical studies in this direction are minimal, undermining the fatality of this lethal disease and possible outbreak.

**Conclusion:** To our knowledge, this is the first bibliometric analysis of Nipah virus-related documents. It urgently demands expedited research in this direction before it is too late.

**Author Summary:** Nipah virus, a rare but deadly bat-borne pathogen, has gained increased attention in recent years. Our pioneering bibliometric analysis reveals a surge in research interest. This study underscores the pressing need for accelerated research on drugs and vaccines to combat this lethal disease and prevent potential outbreaks. Urgency is paramount.

## 1. Introduction

In light of the recent global pandemic caused by the novel coronavirus and its significant economic repercussions, there has been increasing attention on research into viruses with potential risks, one of which is the Nipah virus (NiV). The earliest reports of NiV infecting humans and causing severe health outcomes, such as acute respiratory infections and fatal encephalitis, can be traced back to 1999 in Malaysia (1). Subsequent small-to-moderate outbreaks have been documented in Asian regions, including Bangladesh (2–4) and India (2). In 2018, the World Health Organization (WHO) initially designated NiV as a virus of concern requiring focused attention (5). Following this, organizations like Epidemic Preparedness Innovations (6) and the United Kingdom Vaccine Network (7) have also prioritized NiV for vaccine research and development.

Taxonomically, NiV has been classified as a single-stranded, negative-sense RNA virus, belonging to the genus Henipavirus within the Paramyxoviridae family (8). Its genome consists of a single-stranded, negative-sense RNA of approximately 15kb in length, which is non-segmented **Figure 1A** (3, 9–11). The virus primarily encodes for nucleoprotein (N), phosphoprotein (P), matrix protein (M), fusion glycoprotein (F), attachment glycoprotein (G), and the RNA polymerase or large protein (L). Additionally, the P protein exists in three alternative splicing forms, namely V, W, and C proteins **Figure 1A** (3, 9–11). The F and G glycoproteins on the surface of NiV particles have been found to determine the virus’s invasiveness and control the process of host cell recognition and infection (12, 13). Specifically, the G protein can recognize erythropoietin-producing hepatocellular receptor B2 (Ephrin B2) in either a specific or non-specific manner and facilitate the entry into host cells with the help of the F protein (12, 13). The N and P proteins are primarily responsible for packaging the viral genetic material (14, 15); the M protein is associated with the budding and release of the virus (16); the L protein serves as the virus-specific RNA-dependent RNA polymerase (RdRp) (14); and the variable splicing forms of the P protein—V, W, and C—mediate interactions with the host cell’s interferon response (14, 15).

**Figure 1.**
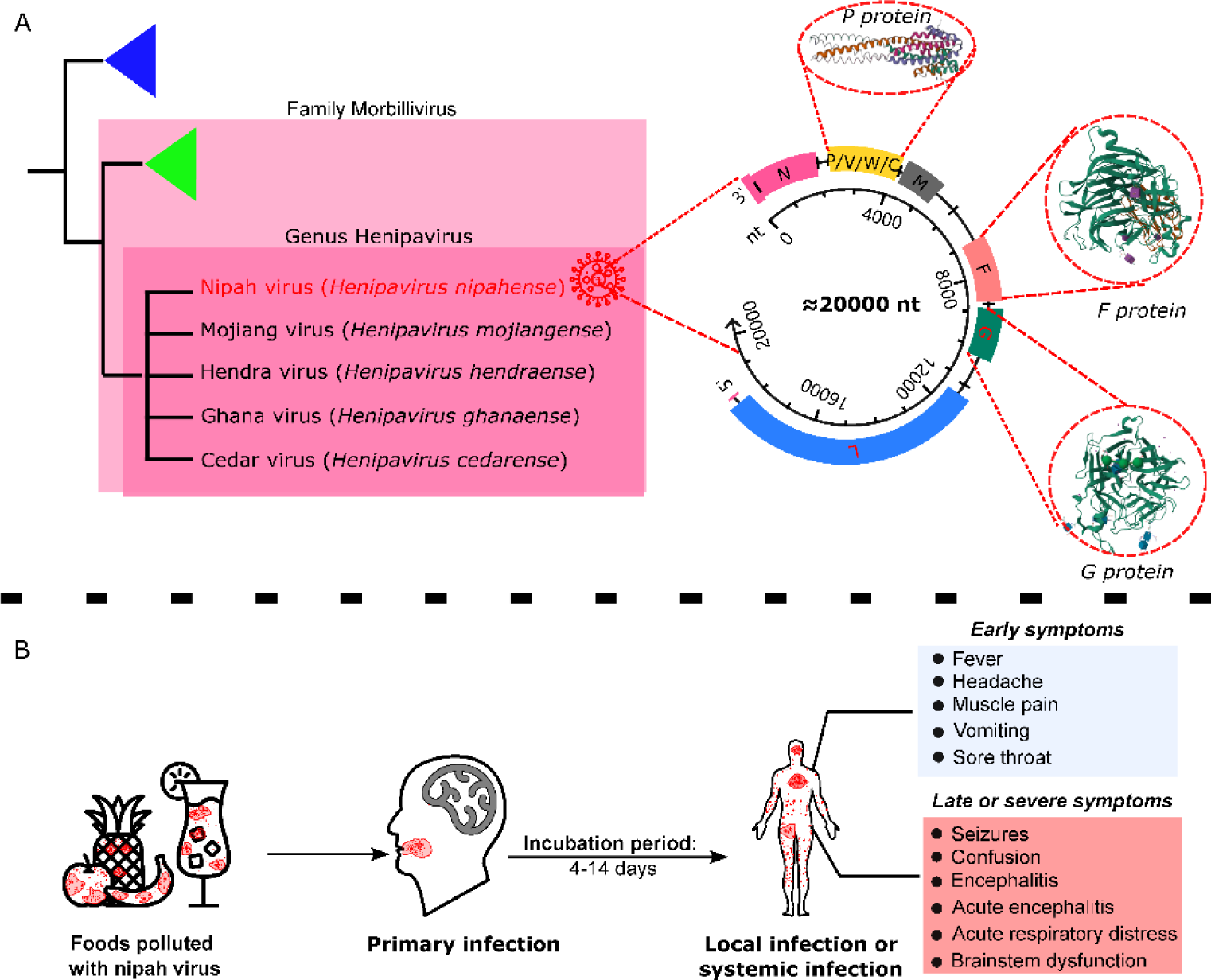
Taxonomy and epidemiological foundation of Nipah virus. Figure 1A illustrates the taxonomy, genome, and principal proteins of the Nipah virus (3, 9–11). The crystal structure of the P protein and its variant spliced forms is derived from the report by Bruhn et al. (4N5B)(15), the crystal structure of the F protein and its complex with Ephrin B2 is from Byrne et al. (8DMJ)(17), and the crystal structure of the G protein and its complex with Ephrin B2 is based on Xu et al. (3D11)(18). Figure 1B presents the infection symptoms and clinical characteristics of the Nipah virus(19–21).

In recent years, there has been a more comprehensive understanding of the infection and transmission mechanisms of the Nipah virus (NiV). The prevailing view posits that NiV is a zoonotic virus, with fruit bats, specifically the species *Pteropus lylei* and *Pteropus hyomelanus*, identified as its primary natural reservoirs(22, 23). Pigs and other mammals act as significant intermediate hosts(22, 23). During outbreaks in Bangladesh and India, the consumption of fruits or fruit products contaminated with NiV emerged as the most likely infection route(22, 23). Furthermore, human-to-human transmission has been confirmed, especially among family members and caregivers of those infected(24). NiV primarily infects the epithelial cells of the respiratory tract and subsequently invades the brain via the olfactory epithelial-neural pathway(19–21). The incubation period generally spans 4-14 days, though it can extend to 40 days in some cases(19–21). Early symptoms often encompass fever, headache, muscle pain, vomiting, and sore throat **Figure 1B** (19–21). These symptoms can progress to dizziness, drowsiness, confusion, brainstem dysfunction, and acute encephalitis. In severe cases, individuals might experience acute respiratory distress, further encephalitis, and seizures **Figure 1B** (19–21). The estimated fatality rate for NiV infections ranges from 40% to 75%. Notably, about 20% of survivors suffer from neurological after-effects, such as epilepsy and altered personalities(19, 20).

Current evidence underscores that NiV is a highly invasive and contagious RNA virus with a pronounced potential for neurological sequelae and elevated fatality rates. As of now, no NiV vaccine has been validated for human use. This study aims to systematically review existing NiV literature using bibliometric methods, analyzing prevailing trends, research hotspots, and inherent limitations. Additionally, we explore the potential applications and impacts of natural products in NiV prevention and treatment. Our bibliometric analysis not only provides insights into the progression and future directions of NiV research but also offers valuable scholarly evidence to inform upcoming studies on NiV prevention and treatment.

## 2. Results and Discussion

### 2.1 Bibliometric Analysis-PubMed-Based Literature

When PubMed was explored for the literature having “Nipah Virus” in their title, it resulted in 601 results (dated 19.09.2023, publishing period is 1999-2023). It includes 65 review articles, four systematic review articles, one meta-analysis, two clinical trials, and one book. When metadata for these 601 articles have been processed in VOSviewer to assess the co-occurrences of MeSH keywords, there are 947 MeSH terms, and 203 co-occurred in the minimum of 5 publications. The top 20 co-occurred MeSH terms are “Nipah virus,” “humans,” “animals,” “henipavirus infections,” “disease outbreaks,” “Chiroptera,” “swine,” “female,” “*Chlorocebus aethiops*,” “male,” “antibodies, viral,” “Malaysia,” “viral proteins,” “Bangladesh,” “Zoonoses,” “viral envelope proteins,” “cell line,” “encephalitis, viral,” “Paramyxovirinae,” and “Vero cells.” **Figure 2** illustrates the interactive mapping between various co-occurring MeSH terms. In the network mapping, some highlighted countries are Malaysia, Thailand, India, Australia, and Bangladesh. Some other co-occurring MeSH terms are Ephrin-B2, monoclonal antibodies, viral envelope proteins, and nucleocapsid proteins. In one randomized controlled trial, Khan and others have observed that one can prevent bat-sap contact if one covers the sap-producing area with bamboo, dhoincha, jute stick, and polythene skirts (25). As of 19.09.2023, this work is cited in 86 literature (Google Scholar), 57 (Scopus), and 67 (Web of Science). Ng and the team have followed up with nine Nipah virus-infected patients for two years and observed that 88.88% of the subjects had developed psychiatric features like major depressive disorder, personality changes, and chronic fatigue syndrome (26). As of 19.09.2023, this work is cited in 70 literature (Google Scholar), 45 (Scopus), and 35 (Web of Science).

**Figure 2.**
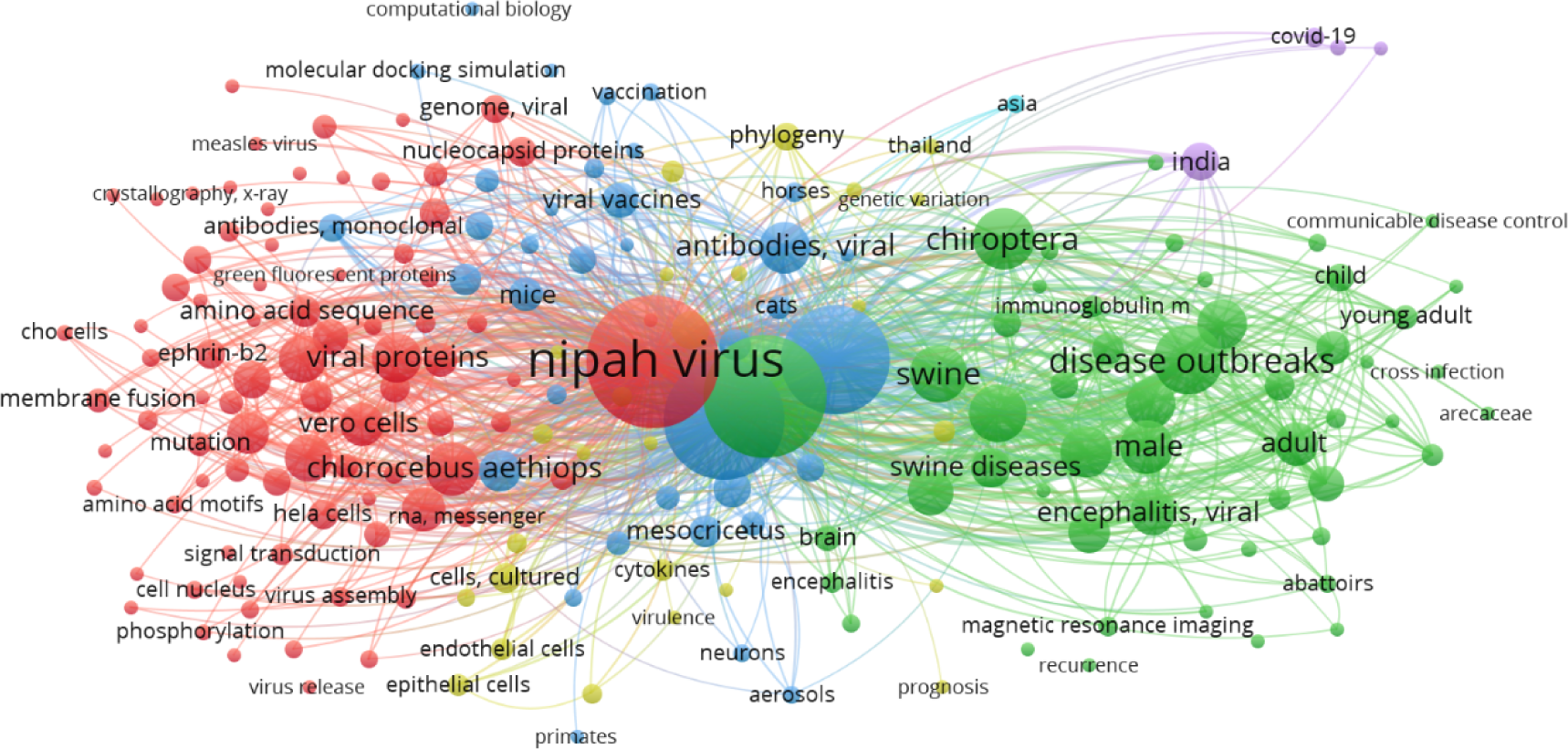
MeSH Term interaction map based on the literature related to “Nipah Virus” (the data obtained from PubMed dated 19.09.2023 and processed in VOSviewer). Distance between two circles will reflect the co-occurrence frequency of those two MeSH terms. More giant the bubble, the more dominant the MeSH term is (27).

### 2.2 Bibliometric Analysis – Literature Retrieved from Web of Science

When Web of Science was explored for articles with “Nipah Virus” in their title, it resulted in 955 literatures (Retrieval date: 19.09.2023; Publication period: 1999-2023). It includes 11 case reports, two clinical trials, 13 books, 66 patents, and 102 review articles. The top 10 highly-cited articles are tabulated in **Table 1**. Each of these ten documents belongs to the type “Article.” It has been observed that Gurley ES, Luby SP, and Broder CC contributed the highest number of publications related to the Nipah virus. “Journal of Virology,” “Emerging Infectious Diseases,” “Journal of Infectious Diseases,” “American Journal of Tropical Medicine and Hygiene,” “International Journal of Infectious Diseases,” and “Plos One” published a maximum number of related publications. In the last five years, there has been exponential growth in the number of publications related to the Nipah Virus, contributing to approximately 35% out of 955 documents. The top five countries to publish the maximum number of publications are the USA (359), Malaysia (111), India (99), Australia (97), and Bangladesh (79). **Figure 3** illustrates the bibliometric data for these 955 documents. **Figure 3A** indicates that the maximum number of publications are visible in 2019, followed by 2020 and 2023. Web of Science has further categorized the documents based on research areas, and **Figure 3B** illustrates that. Most of the documents fall in the category of infectious diseases, followed by virology and microbiology. While evaluating categories in the “research area,” 14 record(s) (1.466%) do not contain data in the field being analyzed. Of 955 documents, 773 are indexed in the Web of Science core collection, while 683 are in Medline (**Figure 3C**). Web of Science has also categorized documents according to “Major concepts.” The majority of the documents fall under the category of infection, followed by “biochemistry and molecular biophysics” and “population studies” (**Figure 3D**). While evaluating categories in “major concepts,” 397 record(s) (41.571%) do not contain data in the field being analyzed. While assessing the citation report generated by Web of Science for these 955 documents, there are 29,077 citations by 7,760 citing articles as of 19.09.2023. The h-index for the Nipah virus was found to be 87 as of 19.09.2023. The top 10 MeSH headings are illustrated in **Figure 4**. While evaluating MeSH headings, 369 record(s) (38.639%) do not contain data in the analyzed field.

**Figure 3:**
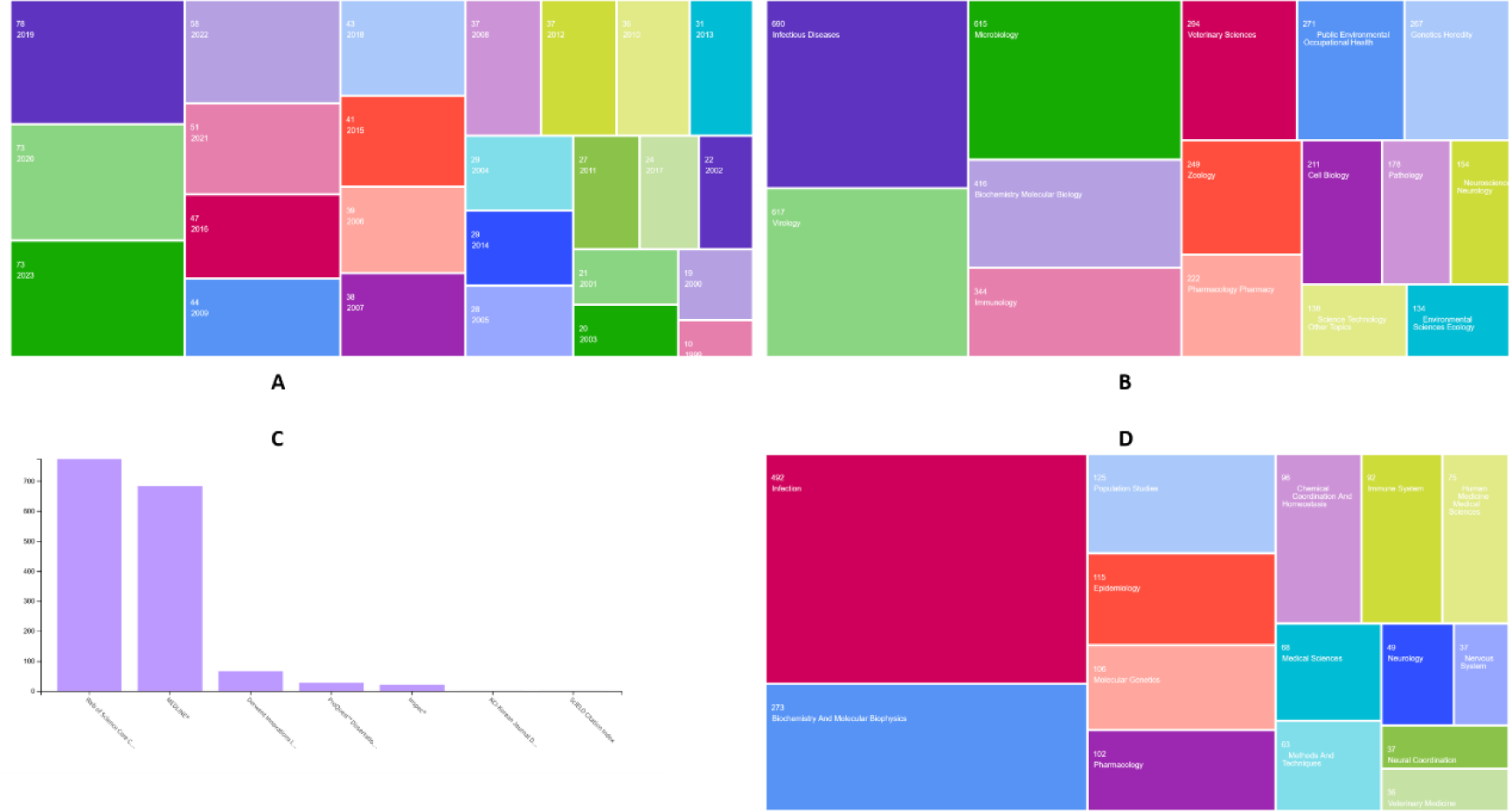
Bibliometric analysis of the “Nipah Virus” related documents, retrieved from Web of Science. A: Publication time; B: Research Area; C: Database covered; D: Major concepts.

**Figure 4:**
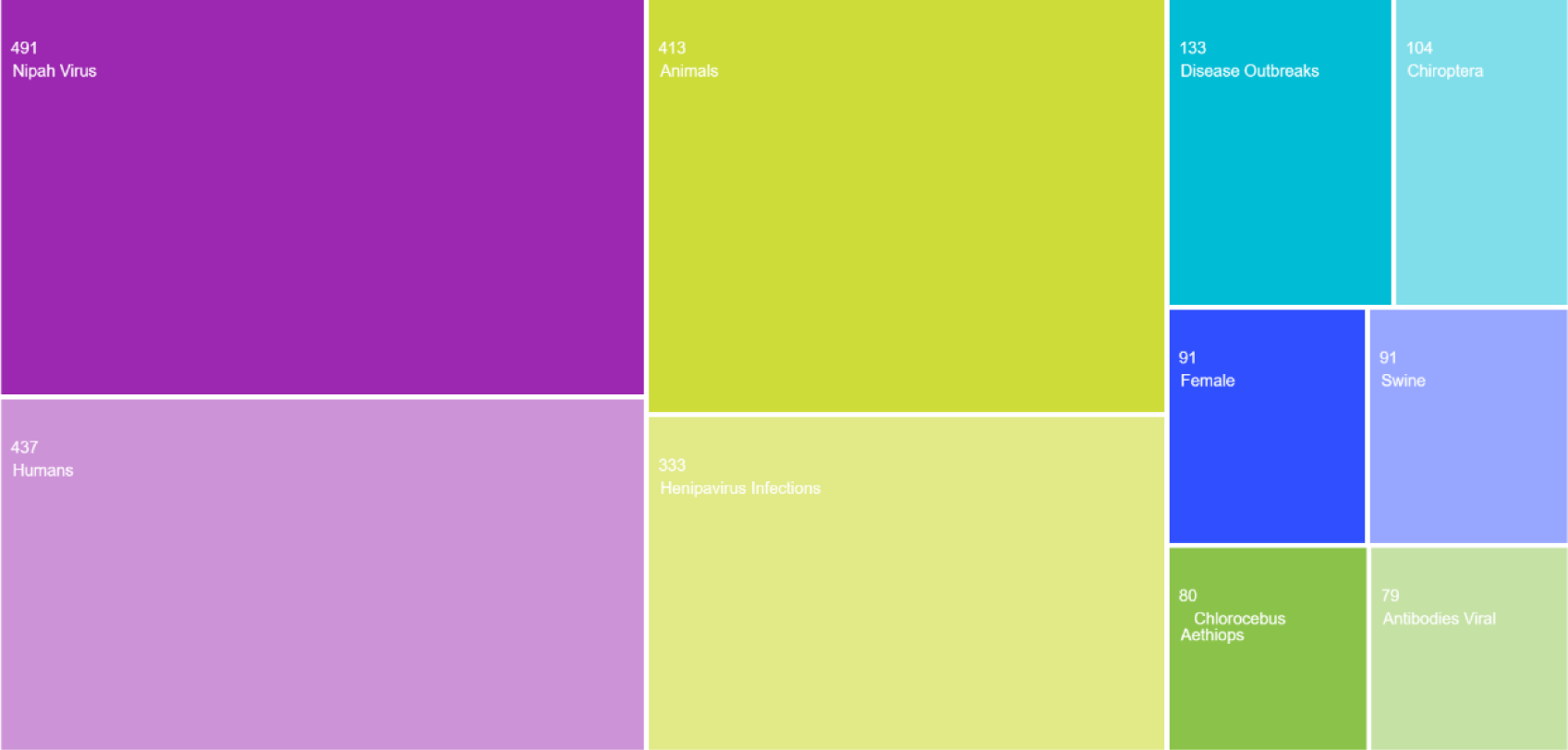
Top 10 MeSH headings in the “Nipah Virus” related documents, retrieved from Web of Science.

**Table 1:**
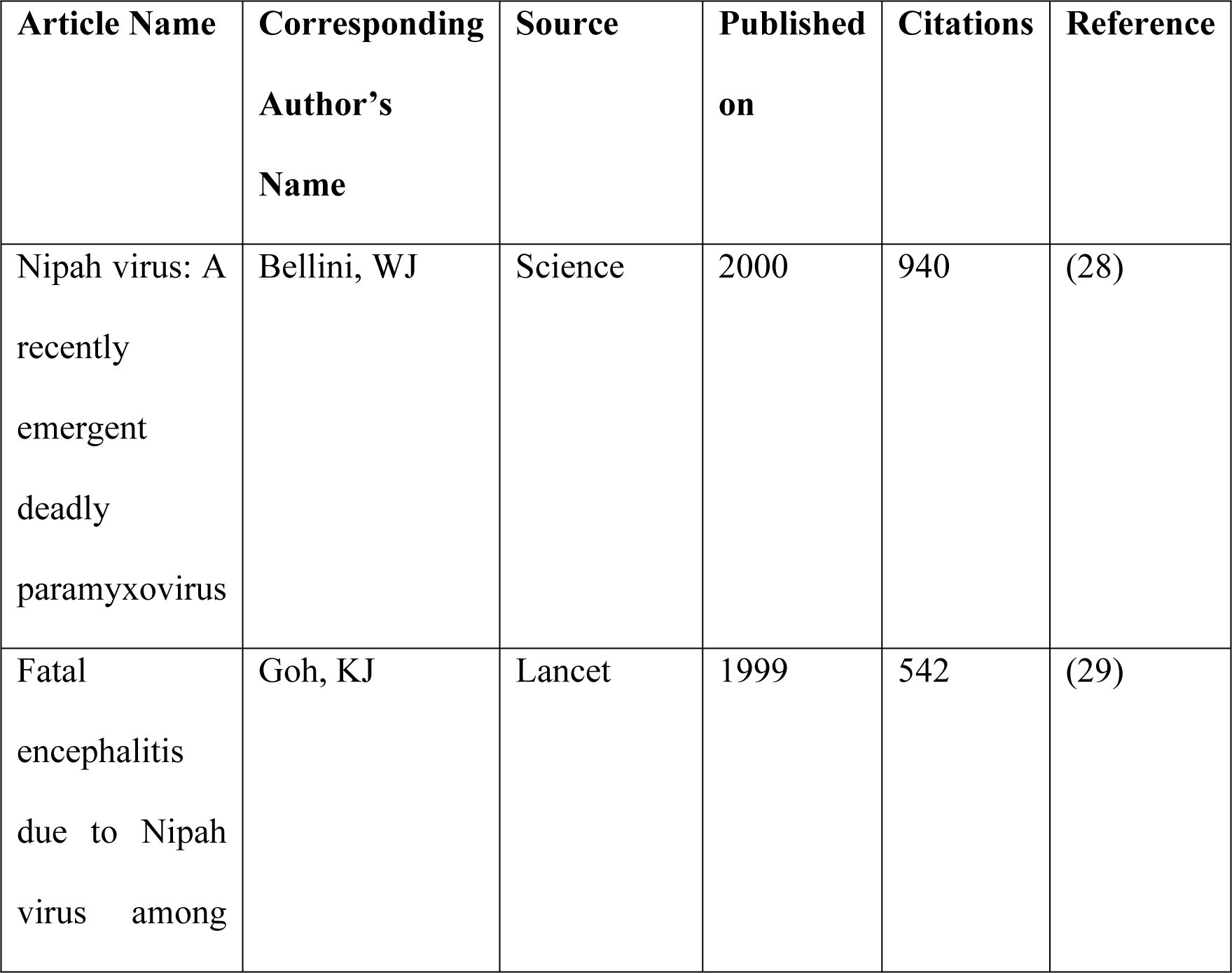
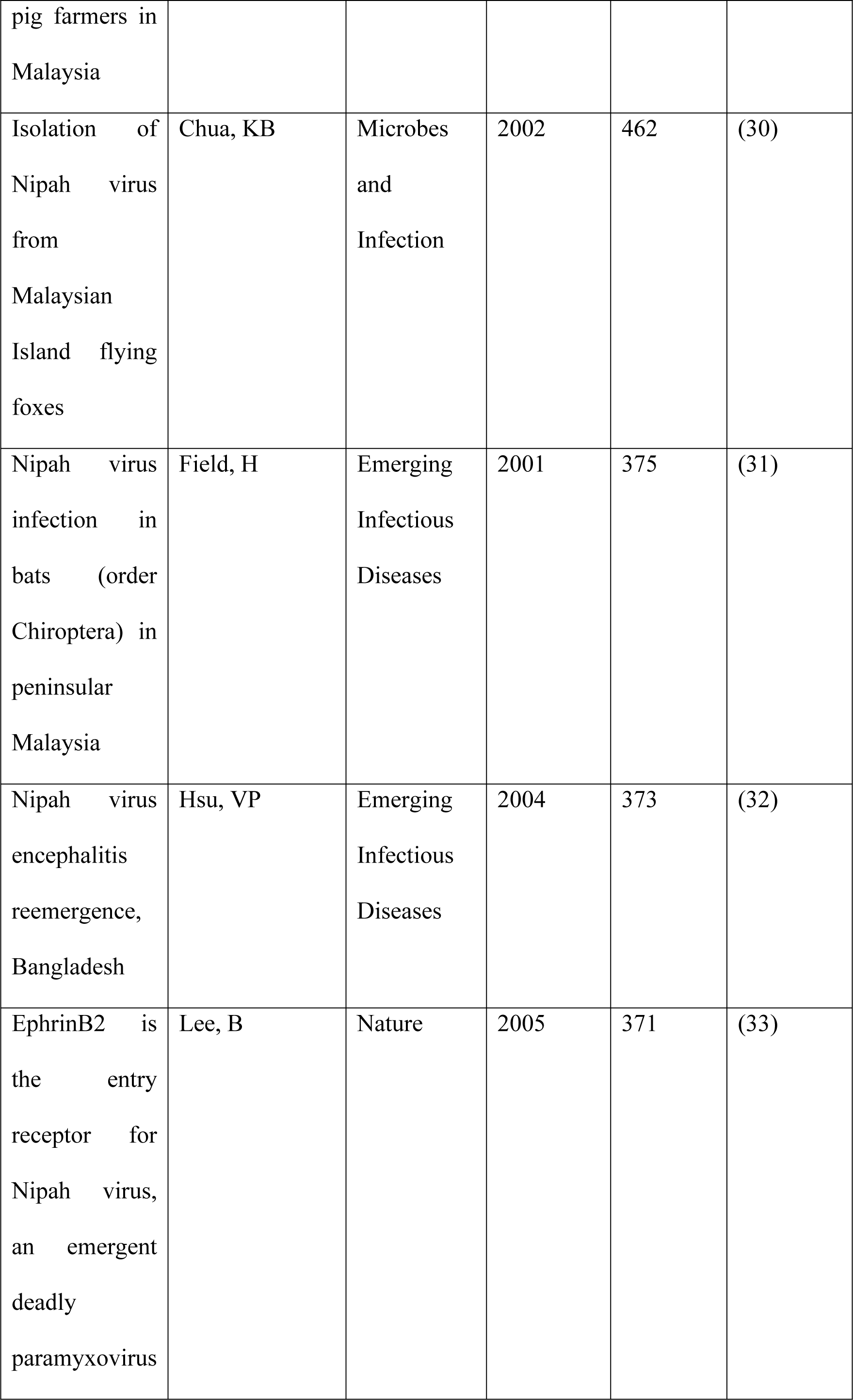
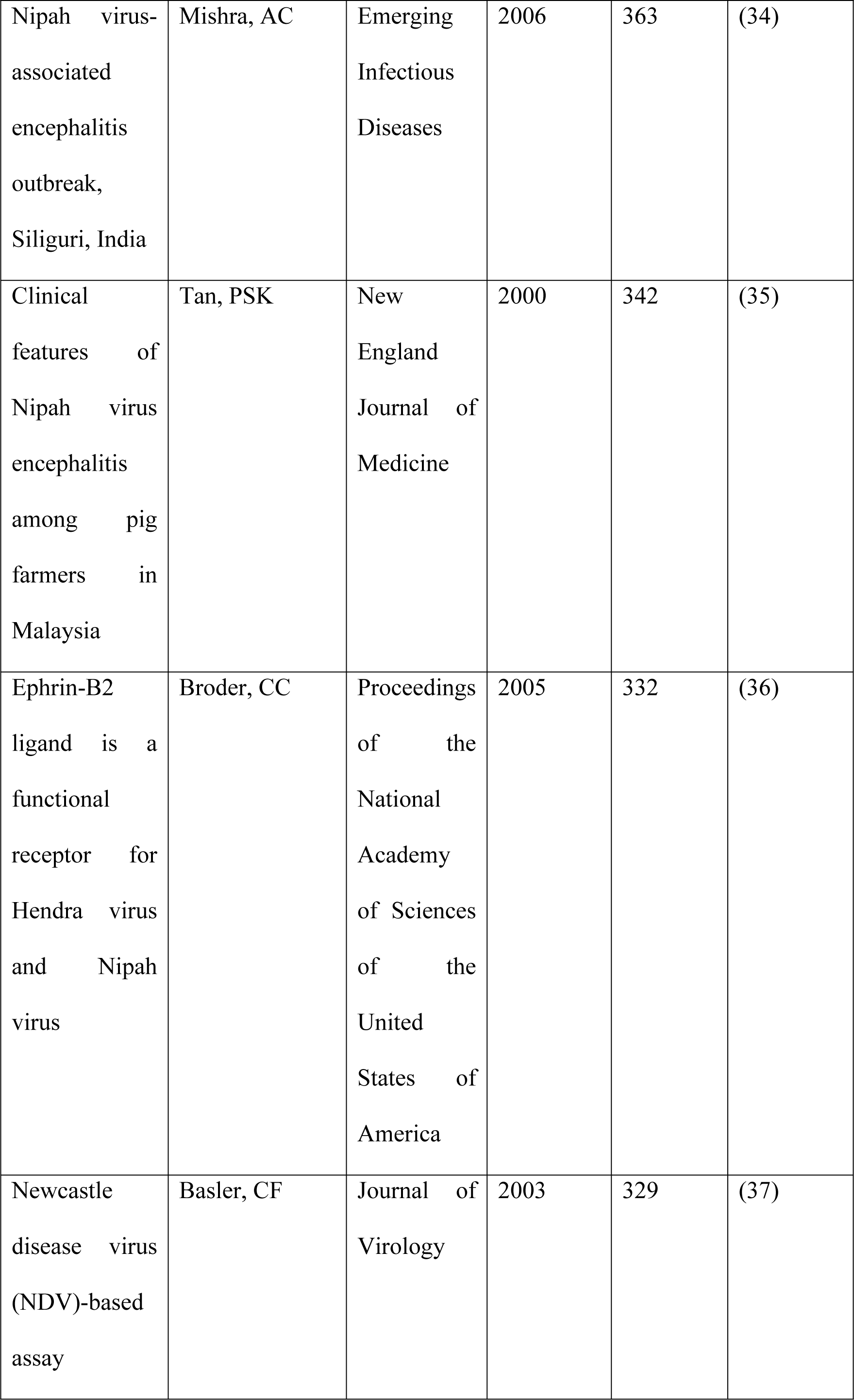
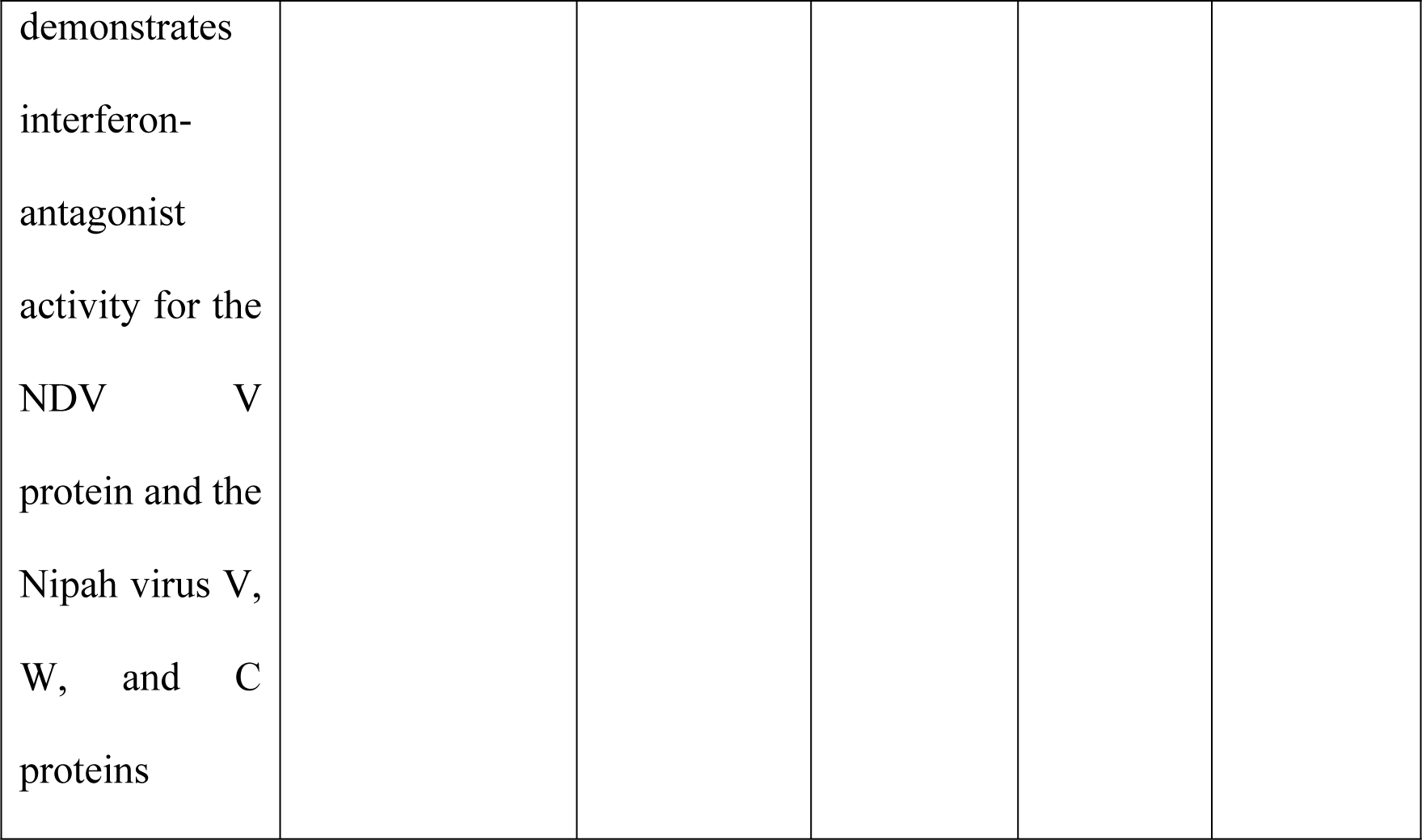
Top 10 highly-cited articles on “Nipah Virus” retrieved from Web of Science on 19.09.2023.

### 2.3 Bibliometric Analysis-Scopus-Based Literature

When Scopus was explored for literature with “Nipah virus” in their titles, it yielded information for 784 documents as of 19.09.2023 (Publication Period 1999-2023). It includes 579 articles, 88 reviews, 29 book chapters, 27 letters, 17 notes, 17 conference papers, 13 editorials, ten erratum, and four short surveys. Just like the observation in Web of Science-based analysis, Gurley ES, Broder CC, and Luby SP contributed the highest number of publications related to the Nipah virus. “Journal of Virology,” “Emerging Infectious Diseases,” “Journal of Infectious Diseases,” “Journal of General Virology,” and “Plos One” published a maximum number of related publications. Though the top five countries contributing publications in this field remain the same, their order and number of publications varied. The top five countries with the maximum number of publications are the USA (341), India (118), Malaysia (106), Australia (85), and Bangladesh (64). Except for India, documents seem to be decreased for the rest of the countries when comparing Scopus with Web of Science. We have used the embedded “Analyze results” tool of Scopus for further analysis. **Figure 5** illustrates the bibliometric analysis of Scopus-retrieved documents having “Nipah virus” in their titles. Contrary to Web of Science, which suggested the maximum number of publications in 2019, Scopus indicated a maximum in 2020. According to the subject area, the maximum number of publications belongs to “Medicine (30.6%),” followed by “Immunology and Microbiology (23.9%),” and “Biochemistry, Genetics, and Molecular Biology (13.6%).” While assessing the citation report generated by Scopus for these 784 documents, there are 30,341 citations as of 20.09.2023. The h-index for the Nipah virus was found to be 90 as of 20.09.2023. Based on this Scopus analysis, the top 10 highly-cited articles are tabulated in **Table 2**. Comparing with the data presented in **Table 1**, we can observe that the top 3 articles remain the same in both tables, with a change in the number of citations.

**Figure 5:**
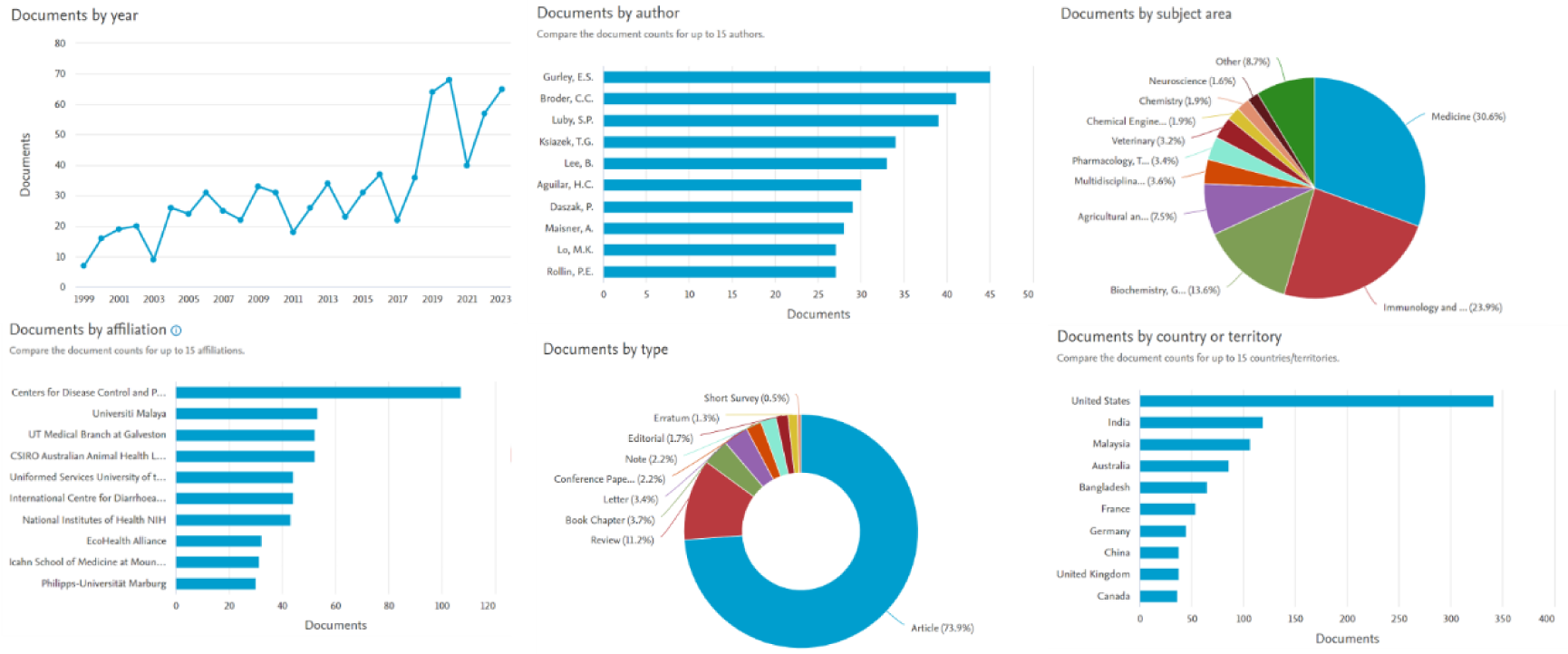
Bibliometric analysis of the “Nipah Virus” related documents retrieved from Scopus.

**Table 2:**
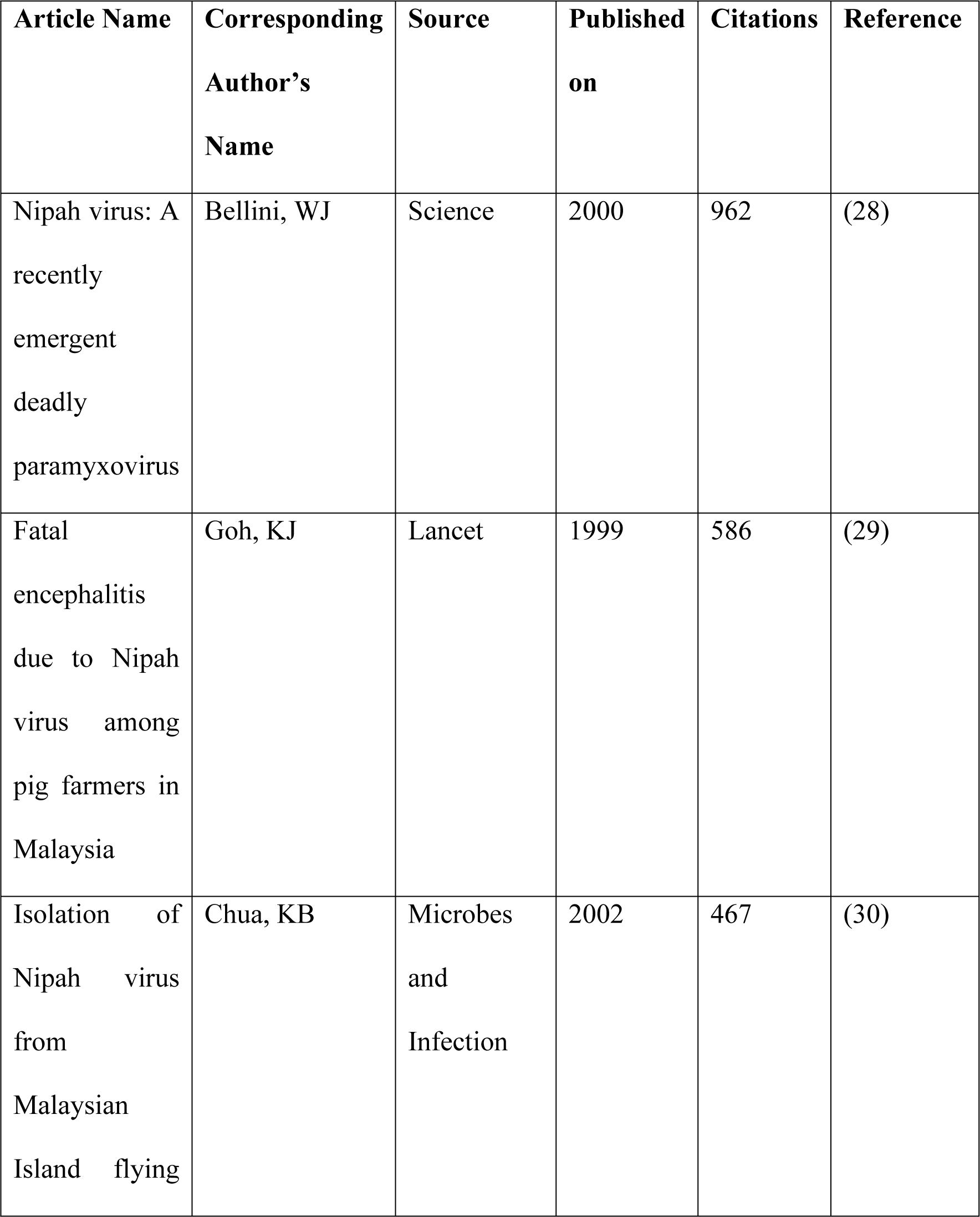
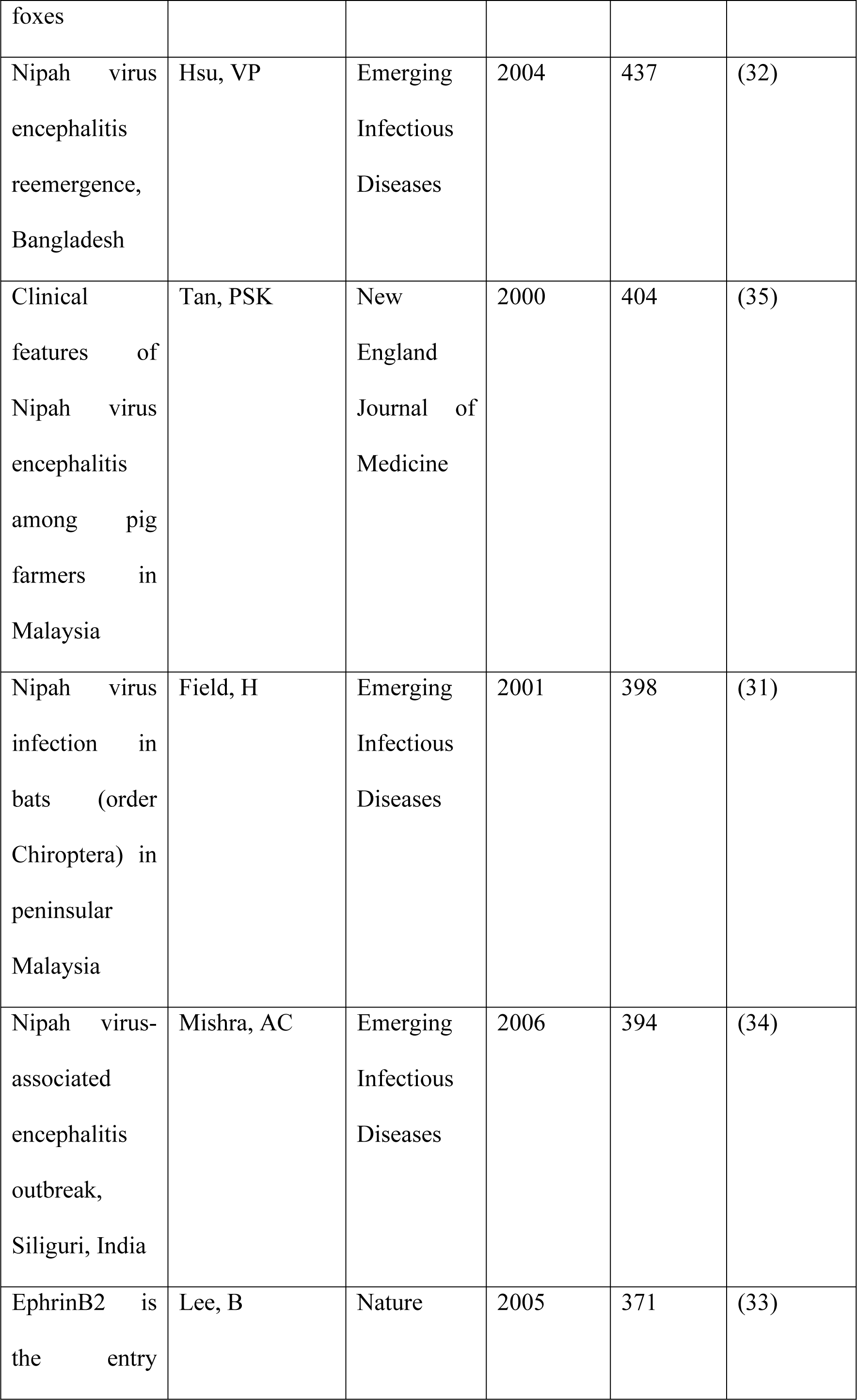
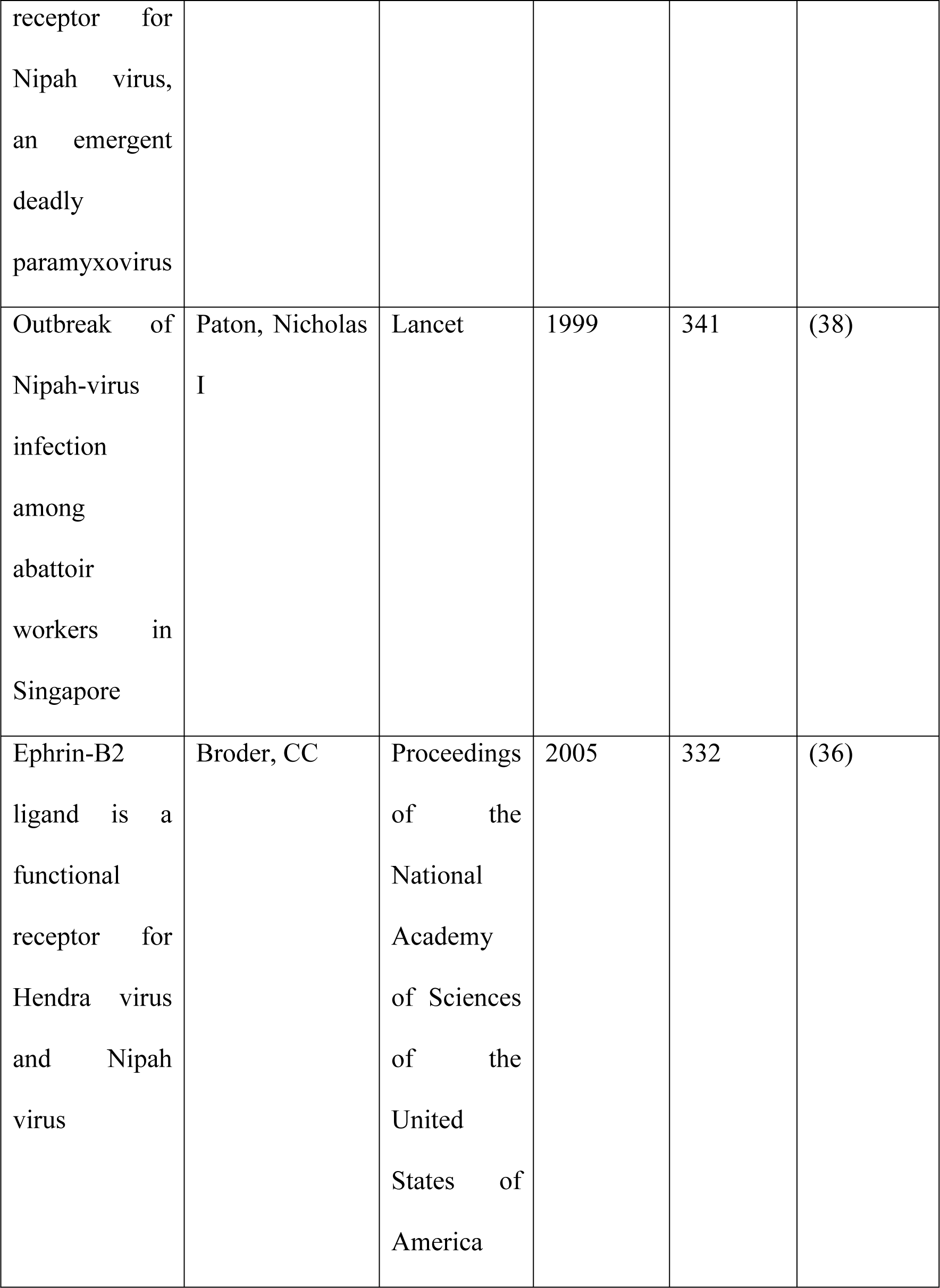
Top 10 highly-cited articles related to “Nipah Virus” retrieved from Scopus as of 20.09.2023.

Metadata from these 784 documents was further processed in VOSviewer for analyzing and seeing trends of index keywords’ co-occurrences. There are 4494 index keywords, and 427 have occurrences in a minimum of ten publications. Some terms like “article,” “humans,” “controlled study,” “priority journal,” “virology,” “animal experiment,” “enzyme linked immunosorbent assay,” “physiology,” “immunology,” “review,” “isolation and purification,” “reverse transcription polymerase chain reaction,” “chemistry,” “amino acid sequence,” “pathology,” “real time polymerase chain reaction,” “signal transduction,” “polymerase chain reaction,” “histopathology,” “protein conformation,” “molecular sequence data,” “clinical article,” “in vitro study,” “western blotting,” “protein protein interaction,” “letter,” “procedures,” “enzyme-linked immunosorbent assay,” “major clinical study,” “nuclear magnetic resonance imaging,” “*in vivo* study,” “health survey,” “flow cytometry,” “geographic distribution,” “neutralization tests,” “sero diagnosis,” “classification,” “sequence analysis,” “electron microscopy,” “protein analysis,” “comparative study,” and “note” have been manually removed from further analysis, as these terms does not seem to provide any insights for future research direction.

The remaining terms were processed for interactive analysis (**Figure 6**). The top ten index keywords are “Nipah virus,” “nonhuman,” “human,” “animals,” “henipavirus infections,” “animal,” “henipavirus infection,” “Nipah virus infection,” “epidemic,” and “female.”

**Figure 6.**
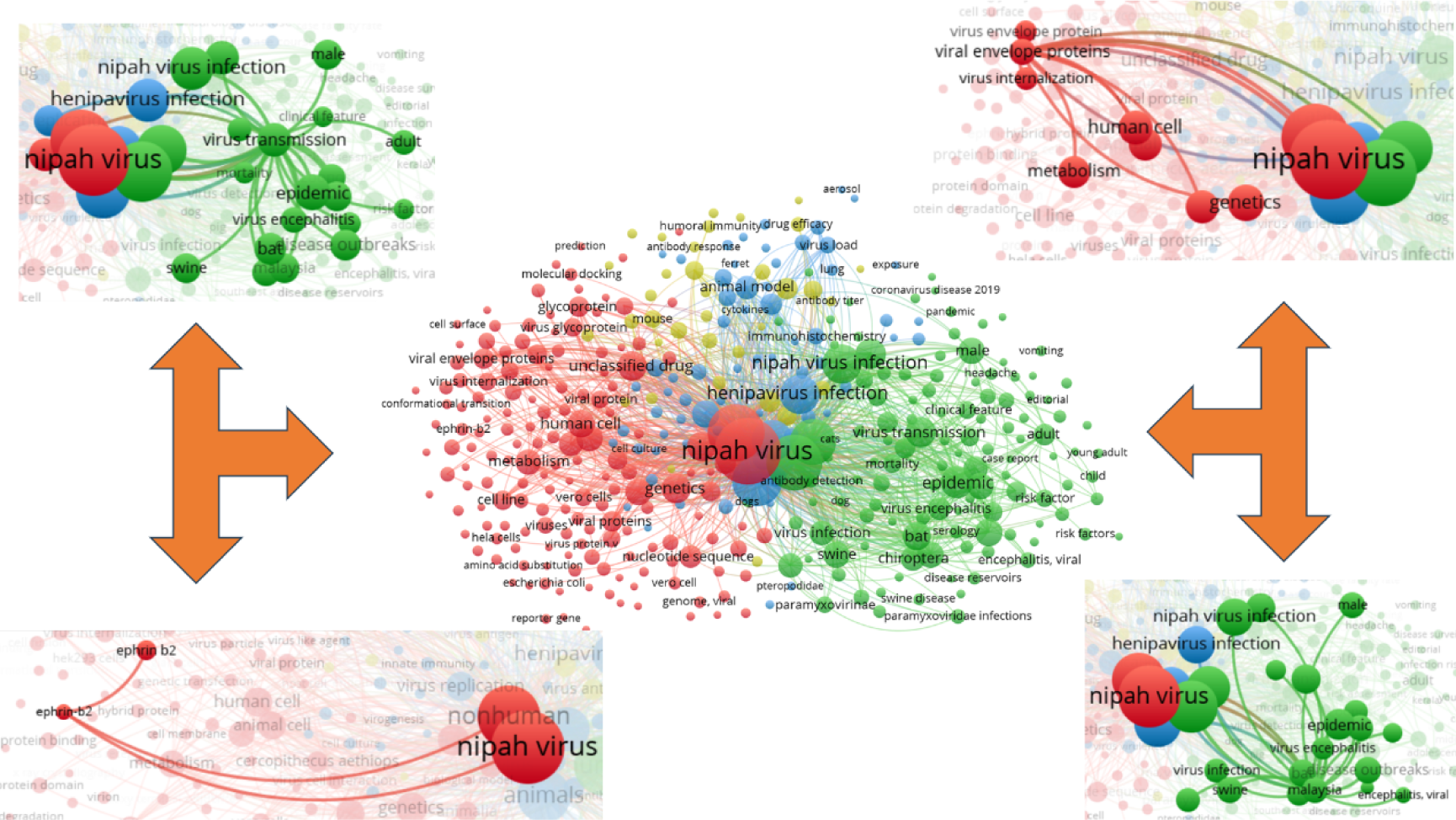
MeSH Term interaction map based on the literature related to “Nipah Virus” (the data obtained from Scopus dated 19.09.2023 and processed in VOSviewer). Distance between two circles will reflect the co-occurrence frequency of those two MeSH terms. More giant the bubble, the more dominant the MeSH term is (27).

## 3. Future Perspectives

### 3.1 Nipah Virus and Targets

As of 20.09.2023, there are 90 documents (excluding review articles) on PubMed that contain “Nipah virus (Title)” and “target (Title/abstract) or receptor (Title/abstract),” which seems to be pretty low in number as compared to the time since it is being first reported. Some of the critical targets and receptors are ephrin-B2 (39, 40), non-structural protein C (41), F protein (42), L protein (43), G glycoprotein (44, 45), nucleocapsid protein (46), V protein (47), P protein (48), W protein (48), and others. Researchers should screen natural and synthetic agents against these targets to find lead compounds.

### 3.2 Nipah Virus and Vaccines

As of 20.09.2023, there are 68 documents (excluding review articles) on PubMed that contain “Nipah virus (Title)” and “vaccine (Title/abstract).” The metadata was processed in VOSviewer for interactive analysis of co-occurring MeSH terms. There are 219 MeSH terms, and 88 have appeared in at least two publications (**Figure 7**). The term “viral vaccines” strongly links with glycoproteins and viral fusion proteins. Foster and others have developed a recombinant vesicular stomatitis virus vectorized vaccine against nonhuman primates (monkeys) (49). Woolsey and the team have also reported a recombinant vesicular stomatitis virus-based vaccine against Nipah virus in African green monkeys, and they have observed long-lasting immunity (50). Keshwara and others proposed a Rabies-based vaccine against the Nipah virus (51). Welch and others have developed a NiVΔF-based vaccine, which protects against lethal Nipah virus infection up to 3 days after vaccination (52).

**Figure 7.**
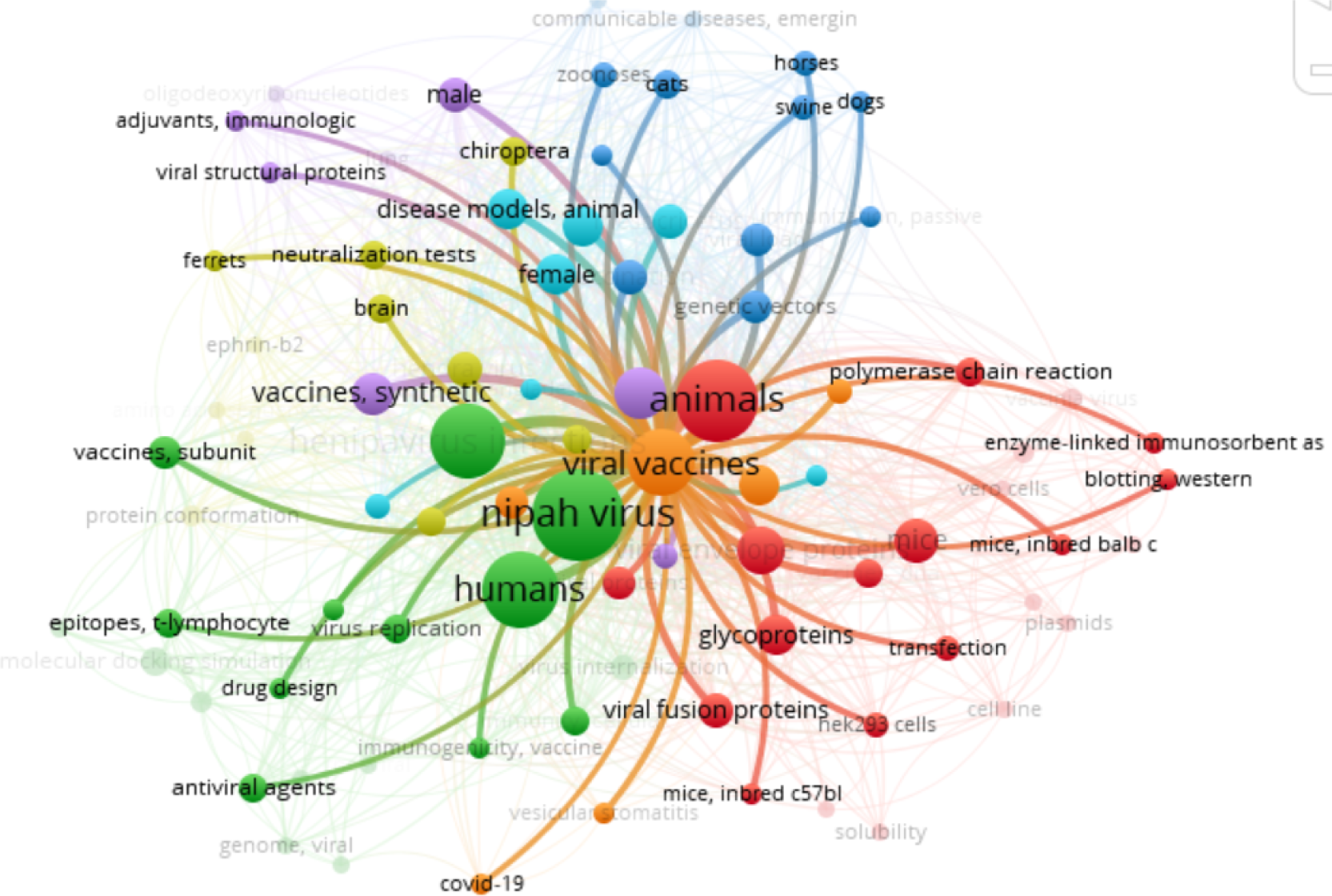
MeSH Term interaction map based on the literature related to “Nipah Virus” and “Vaccine” (the data obtained from PubMed dated 20.09.2023 and processed in VOSviewer). Distance between two circles will reflect the co-occurrence frequency of those two MeSH terms. More giant the bubble, the more dominant the MeSH term is (27).

### 3.3 Nipah Virus and Clinical Studies

While assessing the pattern of clinical studies related to “Nipah virus infection” on https://clinicaltrials.gov/, only four clinical studies are documented there as of 20.09.2023 (**Table 3**). We have observed limitations of scientific shreds of evidence for developing vaccines or drugs against the Nipah virus. Research organizations and funding agencies must direct more energy and funds in this direction.

**Table 3:**
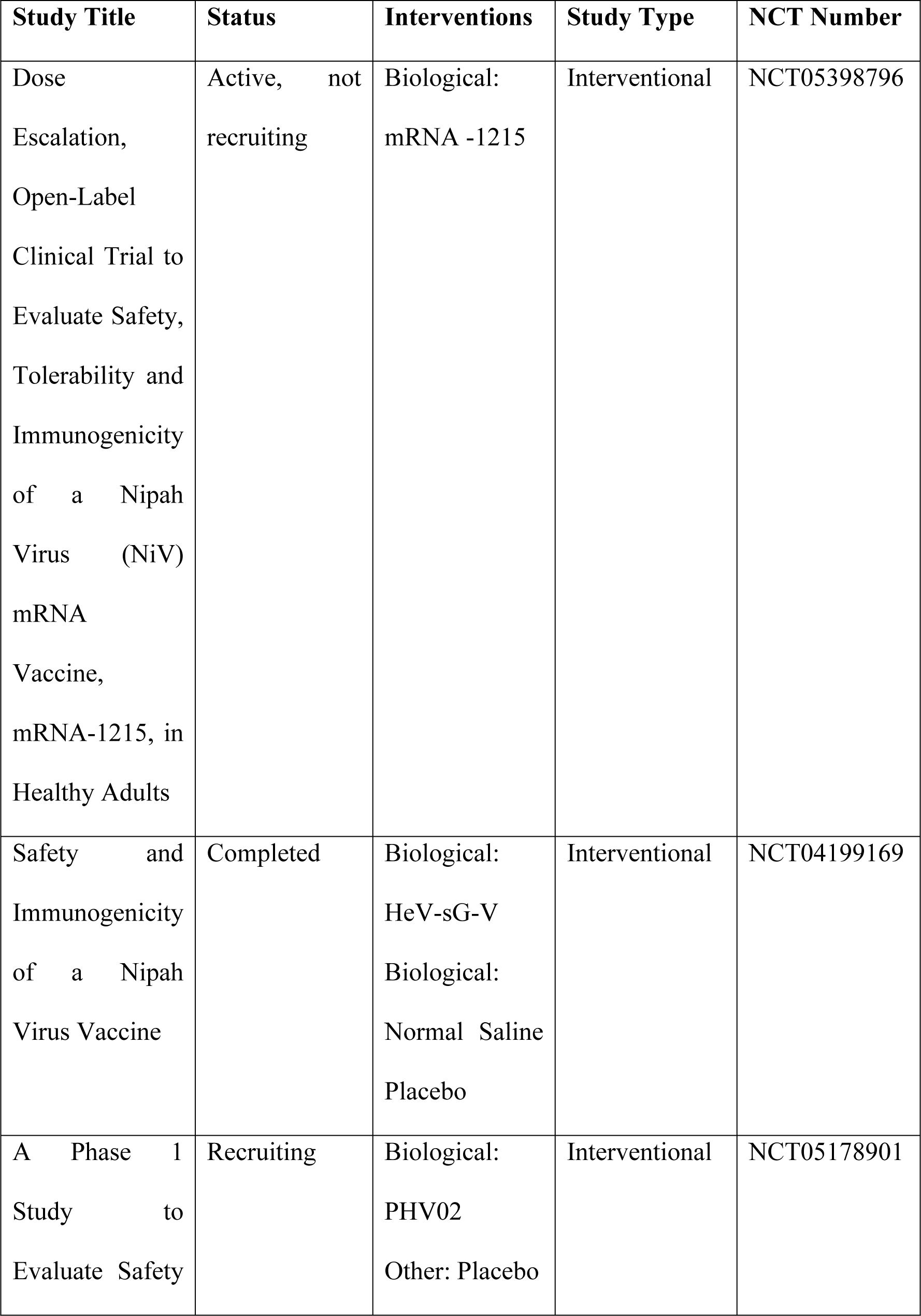
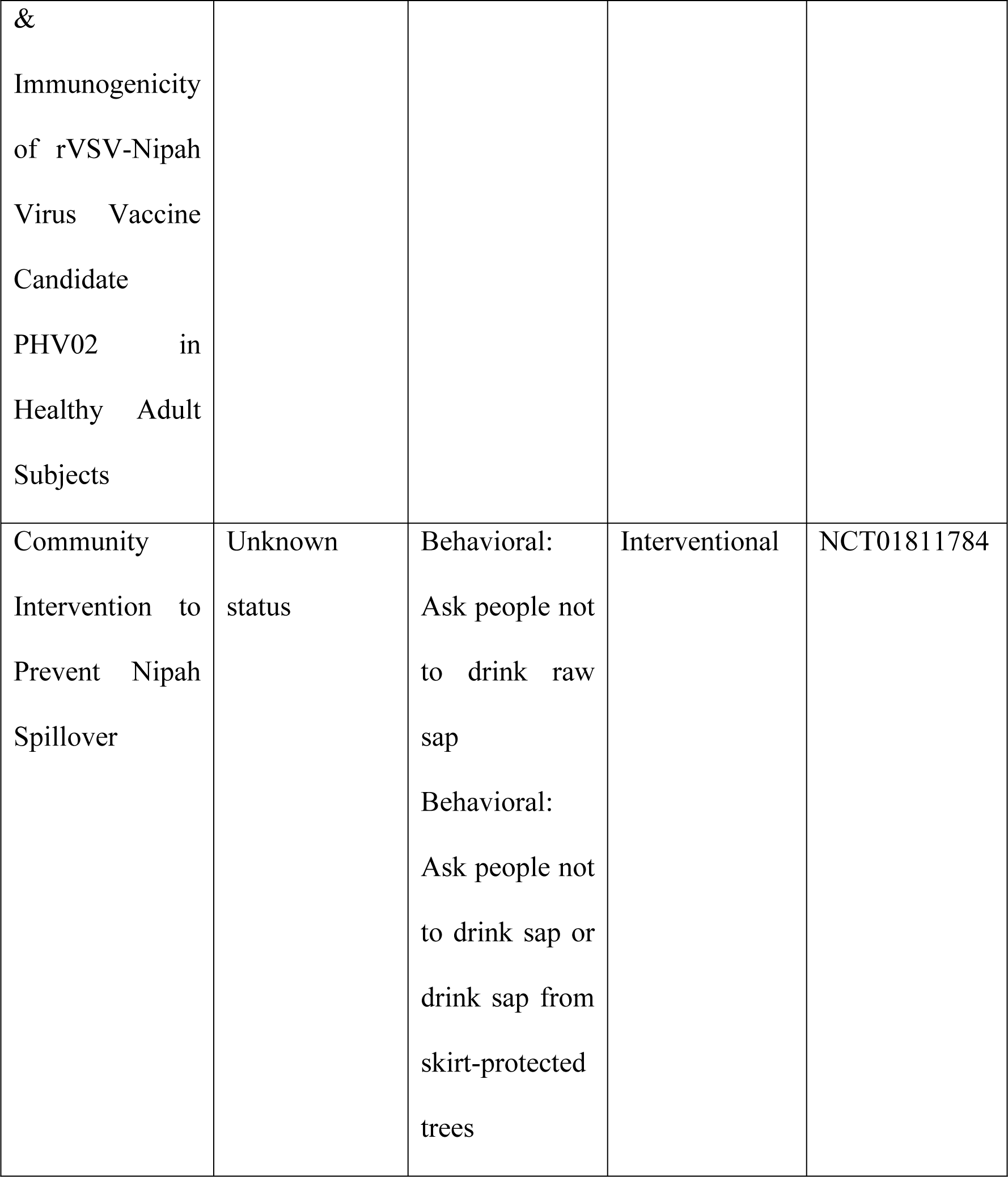
Clinical studies related to Nipah Virus, retrieved from https://clinicaltrials.gov/ on 20.09.2023.

## 4. Materials and Methods

PubMed, Scopus and Web of Science were explored for the documents where “Nipah virus” is in the title. The publication period by default is 1999-2023, and we have not made any restrictions regarding document language. Analysis was made using some in-built tools within PubMed, Scopus, and Web of Science, as well as by processing it in VOSviewer. VOSviewer is a widely used tool for bibliometric analysis (27, 53–55). Clinical studies have been retrieved from https://clinicaltrials.gov/.

## 5. Conclusion

To our knowledge, this is the first attempt to perform bibliometric analysis on the Nipah virus-related documents. The southern state of India (the most populous country in the world), Kerala, has reported patients infected with the Nipah virus in August 2023; two of them have died (56). No drugs and vaccines as of date expose the global population at risk. Because of globalization and urbanization, epidemics are no longer restricted and reserved as epidemics only; opportunistic can cause pandemics like COVID-19 and monkeypox. By putting the up-to-date knowledge about the Nipah virus in this article, we have showcased a niche in this field that could open significant opportunities for clinical and non-clinical researchers. There is an urgent need to explore the translational potential of medicine and vaccines to protect the global community from the Nipah virus.

## Consent for Publication

Not applicable.

## Conflict of Interest

The authors declare no conflicts of interest, financial or otherwise.

## Acknowledgment

None

## Supplementary Materials

None.

## Author Contributions

RKS performed the data curation and analysis. RKS and YZ wrote different manuscript sections. RKS and BS laid the idea, outlined, compiled, and finalized the manuscript. BS arranged the funding source. All the authors have reviewed the final version of the manuscript.

## Funding

This work was supported by the National Natural Science Foundation of China (32070671, 32270690), the COVID-19 research projects of West China Hospital Sichuan University (Grant no. HX-2019-nCoV-057).

## Data Availability Statement

The raw data supporting the results will be available following the proper channel.

## Notes

### Competing Interest Statement

The authors have declared no competing interest.

